# A self-limiting dimeric TIR effector specialised for type III CRISPR-mediated immunity

**DOI:** 10.64898/2026.09.16.751991

**Authors:** Yu Sun, Stephen A McMahon, Silvia Synowsky, Malcolm F White

## Abstract

Antiviral defence systems frequently utilise cyclic nucleotide second messengers. A prominent example is the type III CRISPR-Cas system, which generates cyclic oligoadenylates (cOA) on detecting viral RNA. cOA molecules can bind and activate a wide range of effectors to provide antiviral defence. In both prokaryotes and higher plants, activation of a catalytic Toll/Interleukin Receptor (TIR) domain by multimerization results in degradation of NAD+, limiting cell metabolism and thus viral replication. Here, we describe a CRISPR-associated TIR-containing effector that includes a SAVED (SMODS-associated and fused to various effector domains) domain for nucleotide sensing and a cOA-degrading ring nuclease Crn4 domain. We demonstrate that the TIR-SAVED-Ring nuclease (TSR1) effector binds cA_3_, resulting in activation of the TIR NADase activity. The Crn4 domain, which imposes an unusual dimeric quaternary structure on the effector, degrades cA_3_, providing a mechanism to auto-deactivate the effector. TSR1 is thus a highly unusual example of a dimeric and self-limiting TIR effector in antiviral immunity.

## Introduction

Intracellular cyclic nucleotide signalling is an established paradigm for antiviral defence across all domains of life (1,2). In bacteria, many innate immunity pathways generate cyclic nucleotides in response to phage infection. The Cyclic oligonucleotide-based antiphage signalling system (CBASS), which is ancestral to eukaryotic cGAS/STING, generates cyclic di- and tri-nucleotides (3), whilst Pycsar (pyrimidine cyclase system for antiphage resistance) signals via cyclic mononucleotides (4) and the Thoeris defence system generates cyclic ADP-ribose second messengers (5). All of these innate immunity pathways use cyclic nucleotides as second messengers of infection, amplifying the primary signal and activating a wide range of effector proteins allosterically. Once activated, these effectors typically disrupt cellular metabolism by damaging or degrading key biomolecules, which prevents the completion of the viral replication cycle (2,6).

The most commonly-encountered effector domain in these systems is the TIR (Toll-Interleukin Receptor) domain, which degrades NAD+ upon activation by self-association (reviewed in (2)). Enzymatic TIR domains are found in many bacterial antiphage systems, including type III CRISPR-Cas (7), CBASS (8,9), Thoeris (5), Pycsar (4), Retron (10,11) and Avs (12) defence, as well as many less characterised defence systems (13). Enzymatic TIR domains were first identified and are commonly found in the immune pathways of higher plants (14). A typical mechanism for TIR activation involves self-assembly to complete a shared active site, generally in response to binding a primary or secondary signal of infection (2).

The prokaryotic type III CRISPR-Cas adaptive immune system also functions via cyclic nucleotide signalling. On detection of viral RNA, the catalytic Cas10 subunit generates a range of cyclic oligoadenylate (cOA) species (15). These serve as allosteric activators by binding to the CARF (CRISPR-associated Rossmann Fold) or SAVED domains of a diverse range of effectors (16,17). A smaller number of effectors use alternative nucleotide binding domains; examples include Csx23 (18), CorA (19) and NucC (20,21). CRISPR effectors, once activated, disrupt cellular homeostasis by a wide variety of means including biomolecule degradation and membrane disruption. TIR-domain containing proteins are also associated with type III CRISPR defence: a recent bioinformatic study identified TIR-SAVED effectors in 7 out of ∼1000 type III CRISPR loci (16), while TIR-CARF and SAVED-TIR proteins have also been detected (22,23). Recently, a TIR-CARF effector, designated Cat1, was shown to form filaments on binding cA_4_ in its CARF domain, activating the TIR domain for NAD+ cleavage to provide antiphage defence (7).

Most type III CRISPR systems include a means, designated “ring nuclease” activity, to degrade the cyclic nucleotide activator molecules (24). Some of the effectors have intrinsic ring nuclease activity for auto-deactivation, whilst other type III CRISPR loci encode standalone, extrinsic ring nuclease enzymes. While most of these have a structure related to CARF or SAVED domains (23,24), the recently described Crn4 (CRISPR ring nuclease 4) enzyme uses a completely unrelated fold to bind and degrade a range of cOA species (25). Ring nucleases provide a mechanism to reset the cell once a viral infection has been cleared, or when defence has been triggered aberrantly. This feature seems to be specific to CRISPR systems and has not been observed in the innate immune pathways such as CBASS and Pycsar. This may reflect the fact that these systems tend to be activated later in an infection cycle and often lead to cell dormancy or death (2). Rarely, ring nucleases are fused to CRISPR effectors to generate hybrid enzymes, an example being the Crn1-Csx1 effector (26,27).

Here, we explore the X-ray crystal structure and mechanism of a previously uncharacterised CRISPR effector comprising a fusion of three domains: TIR-SAVED-Ring nuclease Crn4 (hereafter, TSR1) (25). *In vivo*, TSR1 functions as a typical type III CRISPR effector in a plasmid challenge assay. We demonstrate that TSR1 is a dimer in solution and binds cA_3_ sandwiched between two SAVED domains in a head-to-tail conformation, leading to activation of the NADase activity of adjacent TIR domains in a shared active site. The C-terminal Crn4 domain dimerises across the “back” of adjacent TIR-SAVED subunits and functions as a ring nuclease with a preference for cA_3_, providing a mechanism for auto-deactivation. The dimeric organisation of the activated protein contrasts strongly with the larger filaments formed by previously described TIR effectors such as TIR-SAVED (9), Cat1 (7) and TIR-STING (28).

## Materials and Methods

### Cloning and mutagenesis

The synthetic gene (IDT) of *Arachnia propionica* TSR1 (WP_126376302.1) was codon-optimised for expression in *E*.*coli* and cloned into the pEV5hisTEV vector (29), allowing expression with a TEV-cleavable N-terminal polyhistidine tag. The *tsr1* gene was also cloned into Multiple Cloning Site-1 of pRATDuet for the plasmid challenge assay (30). The variants E86Q, H481A, T189E/R190E, N425E/T426E, ΔCrn4 were generated by site directed mutagenesis. The *tsr1* synthetic gene and primer sequences used in this study are listed in Supplementary table S1.

### Protein expression and purification

TSR1 WT and variant proteins were expressed in *E. coli* strain C43 (DE3). Following transformation with the pEV5hisTEV-TSR1 plasmid, a single colony was cultured overnight in 10 mL Luria-Bertani (LB) medium containing 50 mg/mL Kanamycin. The overnight culture was used to seed 1L LB, induced with 0.2 mM IPTG when the OD_600_ reached 0.6–0.8 and incubated at 25 °C overnight with shaking at 200 rpm. Cells were harvested by centrifugation at 5000 rpm, 4 °C for 10 min and stored at -80 °C.

The pellet was lysed by sonication in buffer A (50 mM Tris-HCl, pH 7.5, 0.5 M NaCl, 10 mM imidazole, 10% glycerol) containing protease inhibitors (complete, EDTA-free protease inhibitor cocktail, Roche) and lysozyme at 1 mg/mL final concentration. Cell debris was pelleted by centrifugation at 40000 rpm, 4 °C for 30 min. The cleared lysate was loaded onto a 5 mL HisTrap FF crude column (GE Healthcare) equilibrated with Buffer B (50 mM Tris-HCl, pH 7.5, 0.5 M NaCl, 30 mM imidazole, 10% glycerol).

Unbound proteins were removed by washing in 20CV of Buffer B before TSR1 was eluted in a 30 to 500 mM imidazole concentration-gradient, performed by mixing Buffer B and Buffer C (50 mM Tris-HCl, pH 7.5, 0.5 M NaCl, 500 mM imidazole,10% glycerol). Fractions containing TSR1 were concentrated using a 30-kDa-cutoff Amicon centrifuge filter (Millipore).

To remove the polyhistidine tag, concentrated TSR1 was incubated with TEV protease at a 10:1 (w/w) ratio. The mixture was dialysed overnight at room temperature against Buffer B to remove excess imidazole. Dialysed protein was loaded onto a 5 mL HisTrap FF crude column and the flowthrough collected and loaded onto a size exclusion chromatography (SEC) column 16/60 Superdex 200 (GE Healthcare) equilibrated in Buffer D (20 mM Tris-HCl, pH 7.5, 0.25 M NaCl, 10% glycerol). Fractions from SEC were analysed by SDS-PAGE (NuPage Bis-Tris 4–12%, Invitrogen) with Instant Blue staining (Expedeon). The final concentrated TSR1 was aliquoted, flash frozen in liquid nitrogen and stored at - 80 °C. The concentration of protein was determined by absorbance at 280 nm according to its extinction coefficient. Protein for crystallography was purified in the same manner in buffers excluding glycerol.

### Ring nuclease activity assessed by HPLC

To identify ring nuclease products, 500 nM TSR1 and 50 μM of each cOA species were incubated at 37 °C in buffer containing 50 mM Tris-HCl (pH 8.0), 50 mM KCl and 1 mM EDTA for 1 h. To study the kinetics of TSR1, 250 nM protein and 50 μM cOA species were incubated for 15 min. Reactions were stopped by adding a 2-fold volume of methanol and heating at 95 °C for 10 min, cooling on ice and centrifuging at 12000 rpm for 10 min, 4 °C. The supernatant was transferred to a clean eppendorf tube and dried in a centrifuge concentrator (Eppendorf) at 30 °C for 3 h, then resuspended in 20 μL nuclease-free water. Samples were analysed on a C18 column using a Thermo Ultrapure 3000 chromatography system (Thermo Fisher Scientific). A gradient of Buffer B (Methanol) against Buffer A (20 mM ammonium acetate pH 8.5): 0-0.5 min 1% B, 0.5–6 min 1–15% B, 6–7 min 15–95% B, 7–10 min 95% B, 10–10.5 min 95–1% B, 10.5–15 min 1% B at a flow rate of 0.3 mL/min and column temperature of 40 °C was used. Reaction products were identified by comparing their retention time to standards.

### NADase assay

To analyse the NADase activity of TSR1, εNAD+ (Sigma), a fluorescent NAD+ analogue, was utilised as substrate to be cleaved to εADPR, which generates a fluorescent signal. 40 μL samples containing 0.25 μM protein, 0.5 mM εNAD+ in buffer 50 mM Tris-HCl (pH 8.0), 50 mM KCl and 1 mM EDTA were loaded into 96-well plate (96 well non-binding microplates, Black, F bottom, Greiner). The reaction samples were incubated at 37 °C in a FluoStar Omega microplate reader (BMG Labtech) and the fluorescence signal was collected with filter 300 nm and 410 nm, 15 flashes, 280 cycles with cycle time 20 s. At cycle 50, 0.5 μM cOAs were immediately added to wells to activate the reaction.

### Dynamic light scattering

Dynamic light scattering measurements were performed with the Zetasizer Nano S90 (Malvern) instrument. 1 mg/mL TSR1 was prepared with 1.5-fold excess of cA_3_ in Buffer D.. After centrifugation at 10000 rpm for 10 min at 4 °C, the sample was analysed. Measurements were carried out at 25 °C with 3 measures of 13 runs. The curves of TSR1 and variant with or without cA_3_ are the mean of three technical replicates.

### Analytical size exclusion chromatography

The oligomeric state of TSR1 and its variants was assessed by injecting 100 µL of protein (1 mg/mL) onto a Superose 6 Increase 10/300 GL size-exclusion column (GE Healthcare) using a 100 µL Hamilton syringe. The column was equilibrated with 20 mM Tris-HCl (pH 8.0) containing 250 mM NaCl. To analyse the oligomeric state of TSR1 in the presence of cA_3_, the protein was incubated with a 1.5-fold molar excess of cA_3_ before centrifugation at 12,000 rpm for 10 min at 4 °C. The clarified sample was then loaded onto the size-exclusion column in the same buffer conditions and compared to gel filtration standards (Bio-Rad) consisting of γ-globulin (158 kDa), BSA (66 kDa), ovalbumin (44 kDa) and myoglobin (17 kDa).

### Intact protein mass spectrometry

The protein sample was analysed on a Waters Xevo G2TOF LCMS instrument to determine its molecular weight under denaturing conditions. The protein was diluted to 1µM and 20 µL was injected onto a Waters MassPrep column cartridge. The flow rate was set to 200 μL/min. The gradient was run from 98% A and 2% B to 2% A and 98% B over 4 mins, held for 0.5min, before returning to 98% A and 2% B (A= 95% H_2_O 5% acetonitrile,1% formic acid, B= 5% H_2_O 95% acetonitrile 1% formic acid). MS data were collected from 500-3500 m/z in positive ion mode and scans across the protein peak elution were combined. The raw spectra were processed to mass using the Waters MaxEnt algorithm using a peak width at half maximum (PWHM) of 0.4 Da and a resolution of 0.1 Da. An internal lock mass of Leucine Enkephalin was used. The instrument mass accuracy was checked with a solution of Horse heart myoglobin and was within +/- 1Da.

### Native Mass Spectrometry

Protein samples were buffer exchanged into 150 mM ammonium acetate using Slide-A-Lyzer Mini Dialysis Units (2,000 MWCO; Thermo Fisher Scientific). For ligand-bound samples, proteins were incubated with 0.5x or 3x molar equivalents of cA_3_ on ice for 30 min before centrifugation at 12,000 rpm for 10 min at 4 °C. Native mass spectrometry was performed using protein samples at a final concentration of 5 µM. Analysis of the intact protein complex and subcomplexes was performed using a QToF Ultima (Waters), modified to allow the transmission of large macromolecules (MS Vision). Typical spraying conditions were capillary voltage 3kV and sample cone voltage 200V. The collision energy of the ions was optimised between 50 V to 200 V using an argon gas pressure of ∼2 x 10^−2^ mbar. The mass spectrometer was externally calibrated using aqueous caesium iodide (100 mg/mL) solutions.

### Plasmid challenge assay

The *M. tuberculosis* (Mtb) type III CRISPR system was utilised to recognise the target and subsequently produce cyclic oligoadenylate (cOA) signalling molecules to activate the effector. Recipient cells were generated from *E. coli* C43 (DE3) expressing pMtbCsm1-5 (encoding *M. tuberculosis* Csm1–5) and pCRISPRTetR (encoding Cas6 and crRNA that targets *tetR*), as previously described (31). pRATDuet-TSR1 expressing TSR1 effectors together with a tetracycline-resistance gene were transformed to recipient cells and incubated for 3 h at 37 °C, 210 rpm. Cultures were serially 10-fold diluted from 10^0^ to 10^-3^, and 3 μL of each was applied to LB agar containing 100 μg/ml ampicillin, 50 μg/ml spectinomycin, 12.5 μg/ml tetracycline. In addition, 0.2 % w/v D-lactose and 0.2 % w/v L-arabinose were added to LB agar for full induction of MtbCsm complex and the effectors. Plates were incubated overnight at 37 °C. Each experiment was performed with two independent biological replicates using two technical replicates. Colony-forming units (CFU mL^-1^) were manually counted and calculated by correcting for the dilution factor and plated volume.

### Crystallisation of TSR1

Initial crystallisation conditions for wild-type TSR1 in complex with cA_3_ were identified by sparse matrix screening, where 384 crystallisation conditions were tested on a nanolitre scale. cA_3_ was added to the protein at a 1:1 molar equivalent and incubated at 4 °C for 30 min. Prior to crystallisation, the sample was centrifuged to remove any precipitate. TSR1 – cA_3_ sample and mother liquor (ML) were mixed in 1:1 and 2:1 protein to ML ratios in vapour diffusion sitting drop plates. The experiments were sealed and left to equilibrate at 20 °C. Initial hits produced only poorly formed protein crystals, however, following optimisation diffraction quality crystals grew from 20.5% PEG 3350, 0.05 M potassium sodium tartrate at a protein concentration of 55 mg/ml. Prior to data collection, crystals were cryoprotected with glycerol before cryo-cooling in liquid nitrogen.

### X-ray data processing, structure solution and refinement

X-ray data from TSR1 –cA_3_ crystals were collected at -173 °C on beamline I04 at the Diamond Light Source, using a wavelength of 0.9537 Å. Data were automatically processed using xia2 (32) **/** DIALS (33) to 2.68 Å resolution. The structure was solved by molecular replacement using PhaserMR (34)in the CCP4 suite (35). The search model was generated in AlphaFold3 (36) with initial B-factors modelled in PHENIX (37).

The model was refined via a combination of REFMAC5 (38) and PHENIX with manual model building in COOT (39) between refinement cycles. Electron density corresponding to cA_3_ was clearly visible in the maximum likelihood/σA weighted F_obs_-F_calc_ difference map at 2σ. After several rounds of model building, additional electron density became apparent for ApA>p. Ligand coordinates were generated in ChemDraw (PerkinElmer) where required and restraint libraries were generated using eLBOW in PHENIX, before fitting into the electron density in COOT. Model quality was monitored throughout refinement using MolProbity (40). There are no Ramachandran outliers in the model and fewer than 1% rotamer outliers. Data and refinement statistics are shown in Supplementary Table 2. The coordinates and data have been deposited in the Protein Data Bank with accession code 31JM.

## Results

### TSR1 is active in a plasmid challenge assay

TIR-SAVED effectors are cyclic nucleotide-activated proteins primarily associated with CBASS immunity in bacteria (9). A recent bioinformatic study detected the CRISPR-associated TIR-SAVED-Ring nuclease (TSR1) effector (25). A typical CRISPR gene locus, from *Arachnia propionica*, has TSR1 as the sole predicted ancillary effector (Fig. 1A). To explore the function of TSR1, we first tested the codon-optimised TSR1 effector from *Arachnia propionica* in a plasmid challenge assay established with the *Mycobacterium tuberculosis* type III-A (MtbCsm) CRISPR system (30) (Fig. 1B). Activation of cyclic nucleotide production by the MtbCsm complex in the presence of wt TSR1 resulted in a significant reduction in colony forming units (cfu) (Fig. 1C). Mutation of the conserved catalytic glutamate residue of the TIR domain (E86Q) abolished this activity, implicating the NADase activity in the observed phenotype.

**Figure 1.**
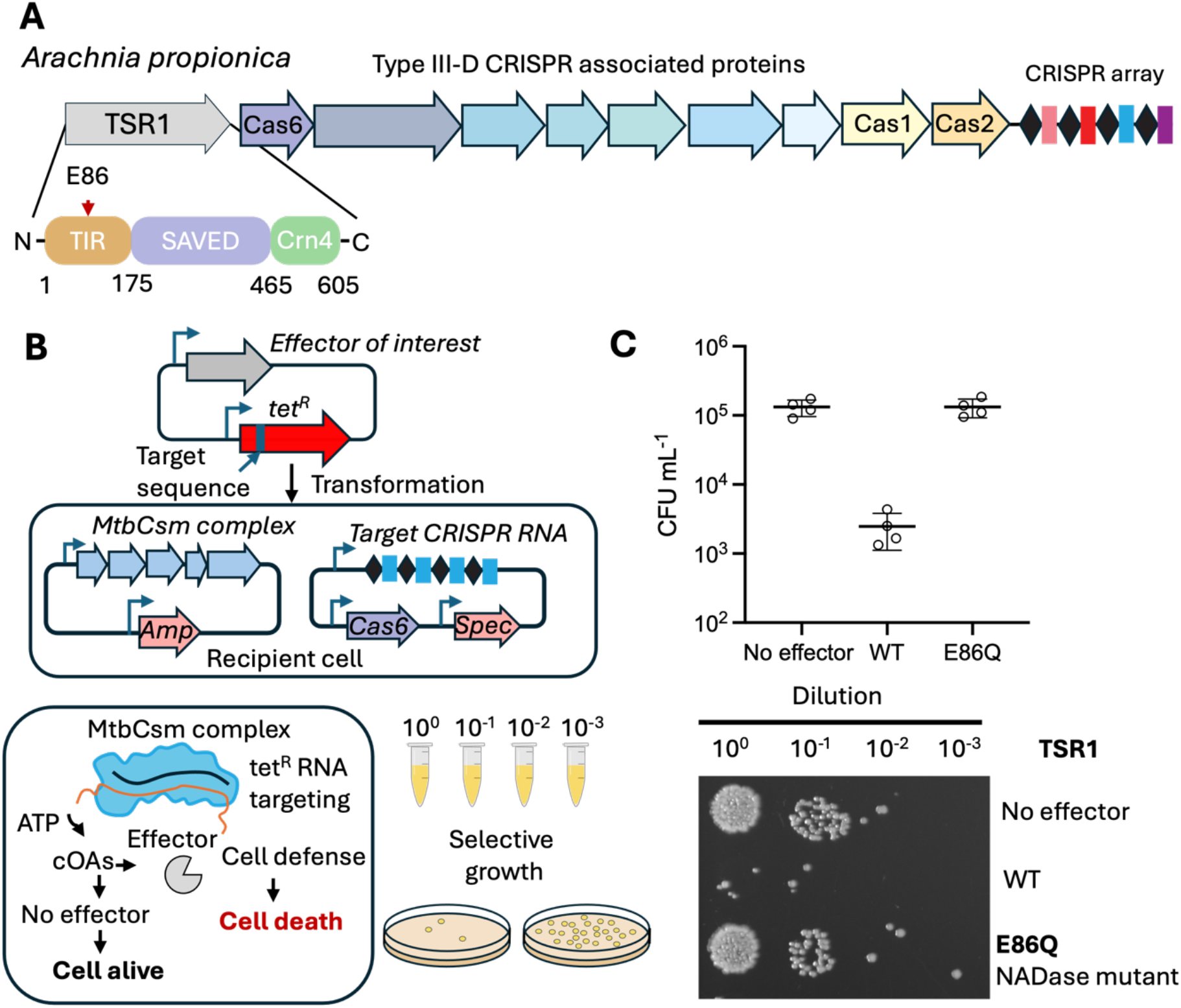
TSR1 functions as a CRISPR-associated NADase effector. **A**. TSR1 (WP_126376302.1) is colocalized with a Type III-D CRISPR locus in the genome of *Arachnia propionica* (NZ_LR134535.1). TSR1 contains a N-terminal TIR domain, central SAVED domain and C terminal Crn4 domain. **B**. Schematic representation of the plasmid challenge assay used to assess effector-mediated immunity. The effector gene is cloned into a plasmid carrying a tetracycline resistance gene and transformed into recipient *E. coli* cells expressing the MtbCsm complex and a crRNA targeting the tetracycline resistance gene. With recognition of the target plasmid, the CRISPR complex synthesizes cOA molecules, which activate the effector. Effector activity is assessed by the resulting growth phenotype of the transformants. **C**. Activation of WT TSR1 by MtbCsm resulted in a significant decrease in colony-forming units (cfu) per ml of culture. The E86Q variant, where the key catalytic residue in the TIR domain is changed, abolishes the phenotype. Other conditions and replicates are shown in Supplementary fig.1.

### The TSR1 TIR domain is activated by cA_3_

To investigate TSR1 at a mechanistic level, we purified the TSR1 protein following expression with a cleavable N-terminal polyhistidine tag in *E. coli*, using immobilised metal affinity and size exclusion chromatography (Supplementary fig. 2). The NADase activity of the purified recombinant protein was investigated using an established fluorogenic assay that detects εADP ribose (9) (fig. 2A). TSR1 was incubated with a range of cyclic nucleotide species, and NADase activity was assessed (fig. 2B), revealing that cyclic tri-adenylate (cA_3_) was the relevant activator (fig. 2B).

**Figure 2.**
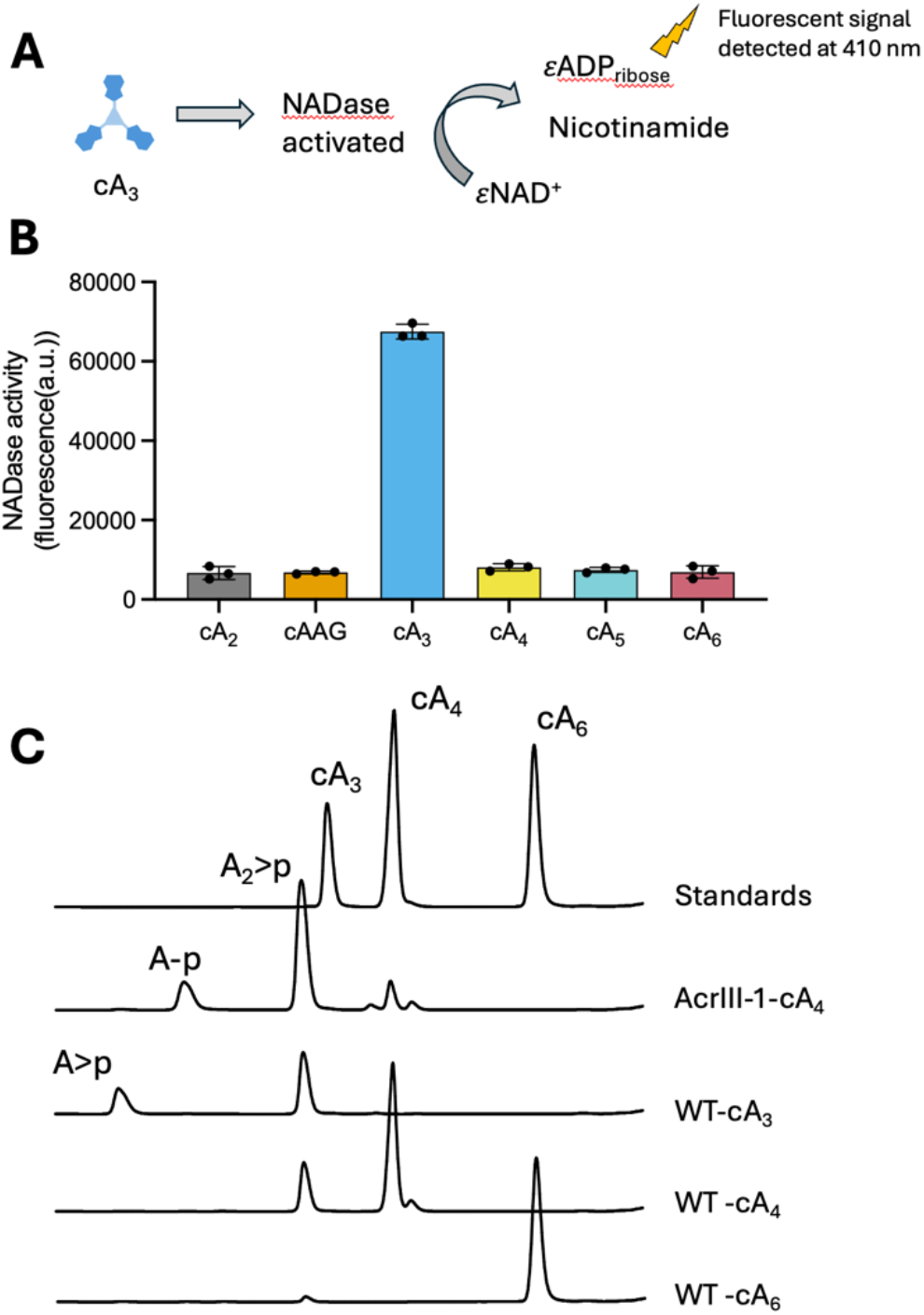
TSR1 is activated by and degrades cA_3_. **A**. Schematic of the NADase assay. **B**. Fluorescence emission was detected at 410 nm upon cleavage of εNAD+ by the TIR domain of TSR1, on incubation with cA_3_. Reactions were incubated at 37 °C with the substrate εNAD+ at 500 µM and WT TSR1 (0.25 µM dimer); 1 µM of the indicated cyclic nucleotide species was added to initiate the reaction. The fluorescence intensity at 90 min was plotted with the standard deviation from triplicate experiments. **C**. HPLC chromatograms of cOA cleavage reactions. Reactions were incubated at 37 °C for 1 h with the indicated cOA species (50 µM) and WT TSR1 (0.5 µM), then analysed by HPLC. The previously characterized ring nuclease AcrIII-1 (42) was incubated with cA_4_ and served as a positive control. Source data are provided as a Source Data file.

### TSR1 degrades cA_3_ and cA_4_

Many CRISPR effectors act as intrinsic ring nucleases, degrading their own activator (24,41). The prediction of a fused Crn4 ring nuclease in the TSR1 structure provided an obvious route for activator degradation. To test this, we incubated TSR1 with cA_3_, cA_4_ or cA_6_ at a dimer:cOA ratio of 1:100, then analysed the reaction products by HPLC (fig. 2C). By comparison with standards, we observed complete degradation of cA_3_ in the 1 h reaction to A_2_ >p and A>p products (where “>” symbolises a cyclic 2’3’ phosphate terminus). The cA_4_ species was partly converted to A_2_ >p under the same conditions (fig. 2C).

### The ring nuclease domain of TSR1 limits the activity of the effector

To investigate the relationship between the ring nuclease and TIR-SAVED domains of TSR1 we generated a H481A variant of TSR1, knocking out the key catalytic residue essential for Crn4-dependent ring nuclease activity (25) (fig. 3A). This variant was purified as for the wild-type protein, but lacked ring nuclease activity *in vitro*, as expected (Supplementary fig. 3). By comparing the WT and H481A variant of TSR1 in the cA_3_-activated NADase assay, we observed a moderate decrease in the activity of the enzyme when the ring nuclease was active (fig. 3B). To investigate this further, we pre-incubated TSR1 with cA_3_ for 40 min before initiating the NADase assay, observing a marked decrease in activity when the ring nuclease was active (fig. 3C). These data are consistent with a role for the Crn4 ring nuclease in limiting the activity of the fused TIR-SAVED effector. Notably, the H481A variant displayed no ring nuclease activity *in vitro*, ruling out a supplementary role for the SAVED domain as a ring nuclease (Supplementary fig. 3B).

**Figure 3.**
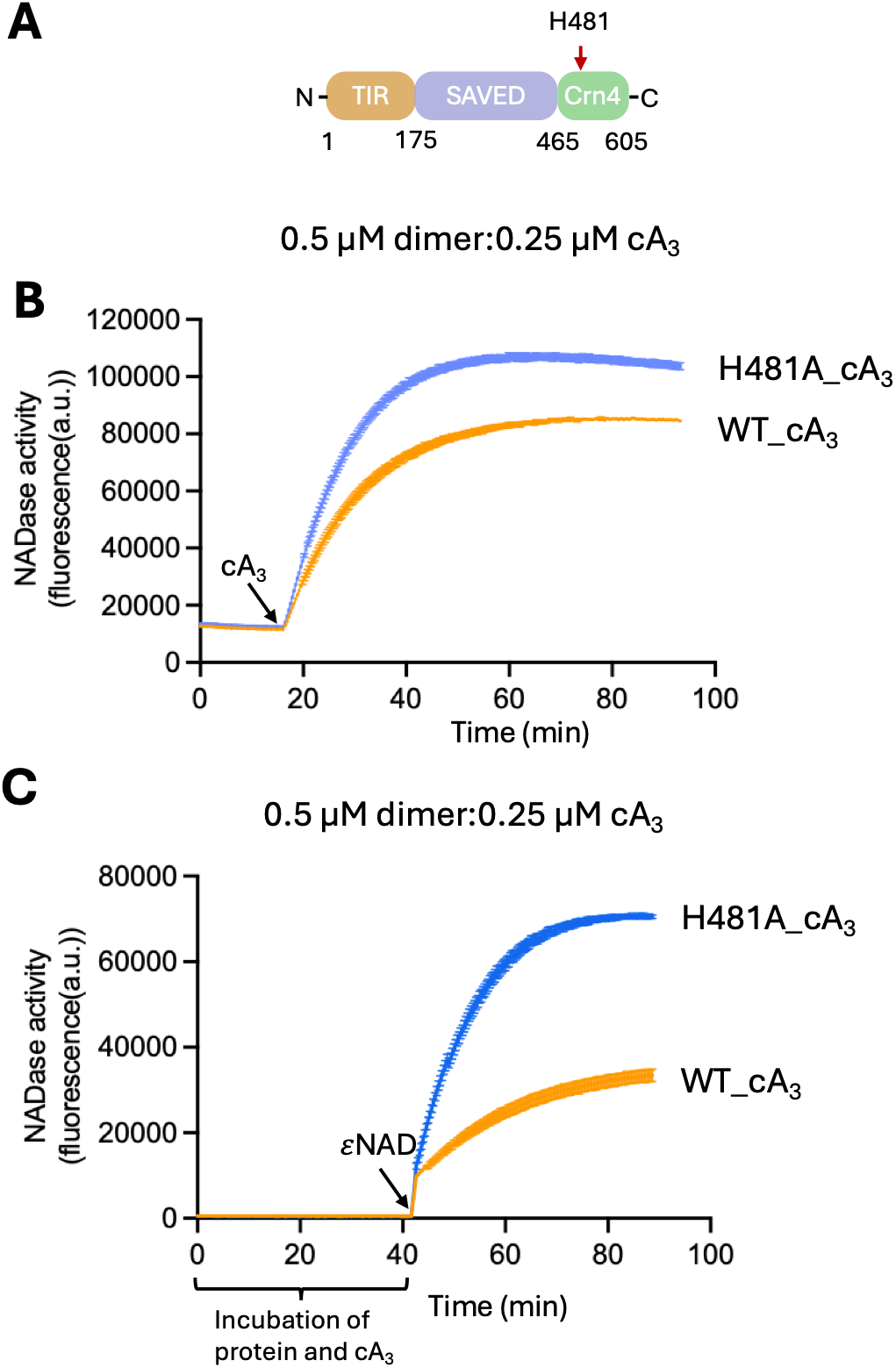
TSR1 is auto-deactivated by the Crn4 domain. **A**. Schematic representation of the H481A variant. **B**. NADase activity of WT and H481A were analyzed by continuous fluorescence assay. Reactions were incubated at 37 °C with 0.5 µM TSR1. 0.25 µM cA_3_ was added at the timepoint indicated to initiate the reaction (arrow). **C**. In this reaction, the WT or H481A variant of TSR1 was pre-incubated with cA_3_ for 40 min before initiating the reaction with εNAD. The plots show means and standard deviations for triplicate experiments.

### Structure of TIR-SAVED-Crn4

To elucidate the molecular structure of the TSR1 effector, we crystallised TSR1 in the presence of cA_3_ and solved the structure at a resolution of 2.68 Å. The crystal structure revealed a dimeric organisation with two TSR1 monomers arranged in a head-to-tail configuration (Fig. 4A), sandwiching a single molecule of cA_3_ in a shared binding site formed by the two SAVED domains. This part of the structure is reminiscent of other SAVED protein complexes bound to cA_3_ or cA_4_, including TIR-SAVED (9), AcrIII-2 (43) and CalpL (44). The TSR1 dimer organises its SAVED and TIR domains in a manner highly reminiscent of two consecutive subunits of the previously studied CBASS TIR-SAVED effector (9) (Fig. 4B). The C-terminal Crn4 domain (Fig. 4C) forms a canonical dimer highly reminiscent of the structure of stand-alone ring nuclease Crn4 (25) (Fig. 4D). The major degradation product of cA_3_, ApA>p, could be modelled into density present in the active site of the Crn4 domain (Fig. 4C).

**Figure 4.**
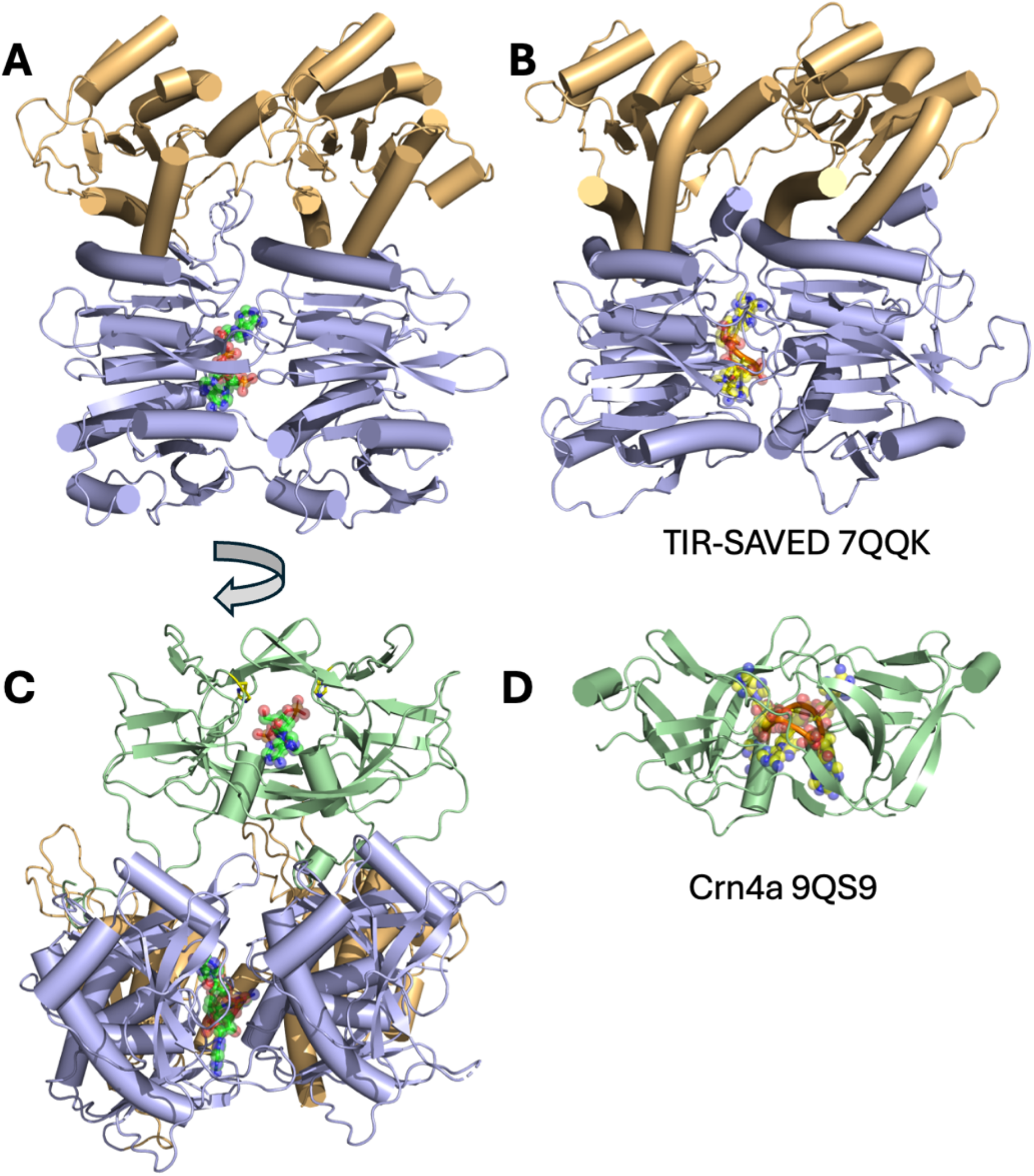
Structure of TSR1 dimer bound to cA_3_. **A**. TSR1 dimeric structure showing the TIR domains in gold and SAVED domains in blue (Crn4 domains omitted for clarity). cA_3_ bound in the interface between SAVED monomers is shown in ball-and-stick representation with atomic colouring. **B**. structure of two adjacent *Microbacterium ketosireducens* TIR-SAVED domains of the active, filament structure formed on cA_3_ binding (PDB 7QQK) (9), coloured as in A. **C**. orthogonal view of the TSR1 dimer rotated by 90 ° with respect to view A. The dimeric Crn4 domain is shown in light green with the ApA>p reaction product in ball-and-stick representation with atomic colouring. **D**. structure of the dimeric ring nuclease Crn4a bound to cA_6_(PDB 9QS9) (25).

### Domain comparisons and nucleotide binding sites

The TSR1 dimeric structure brings the two adjacent TIR domains into close proximity, forming a shared NADase active site, corresponding closely to the structure of the active form of the TIR-containing, cA_4_-activated Cat1 effector (7) (Fig. 5A). In the Cat1 structure, the NAD analogue BAD (benzamide adenine dinucleotide) was used to locate the nucleotide binding site between the two adjacent subunits (Fig. 5A). Turning to the SAVED domains, cA_3_ is well defined at the interface between SAVED subunits (Fig 5B), bound in a planar conformation similar to that observed in the TIR-SAVED complex (9). The conserved residues T189/R190 and N425/T426 are well positioned to provide important interactions with the cA_3_ molecule on either side of the binding site. We mutated these residues in pairs to glutamate, generating the T189E/R190E and N425E/T426E variants, which were no longer functional *in vitro* or *in vivo* (Supplementary fig. 1,4). It should be noted that, due to the head-to-tail conformation of TSR1 monomers, the SAVED domains have externally-facing half-sites for cA_3_ binding (Fig. 5B). In the *M. ketosireducens* TIR-SAVED structure, these sites all participate in binding cA_3_ to stabilise the extended filament (9).

**Figure 5.**
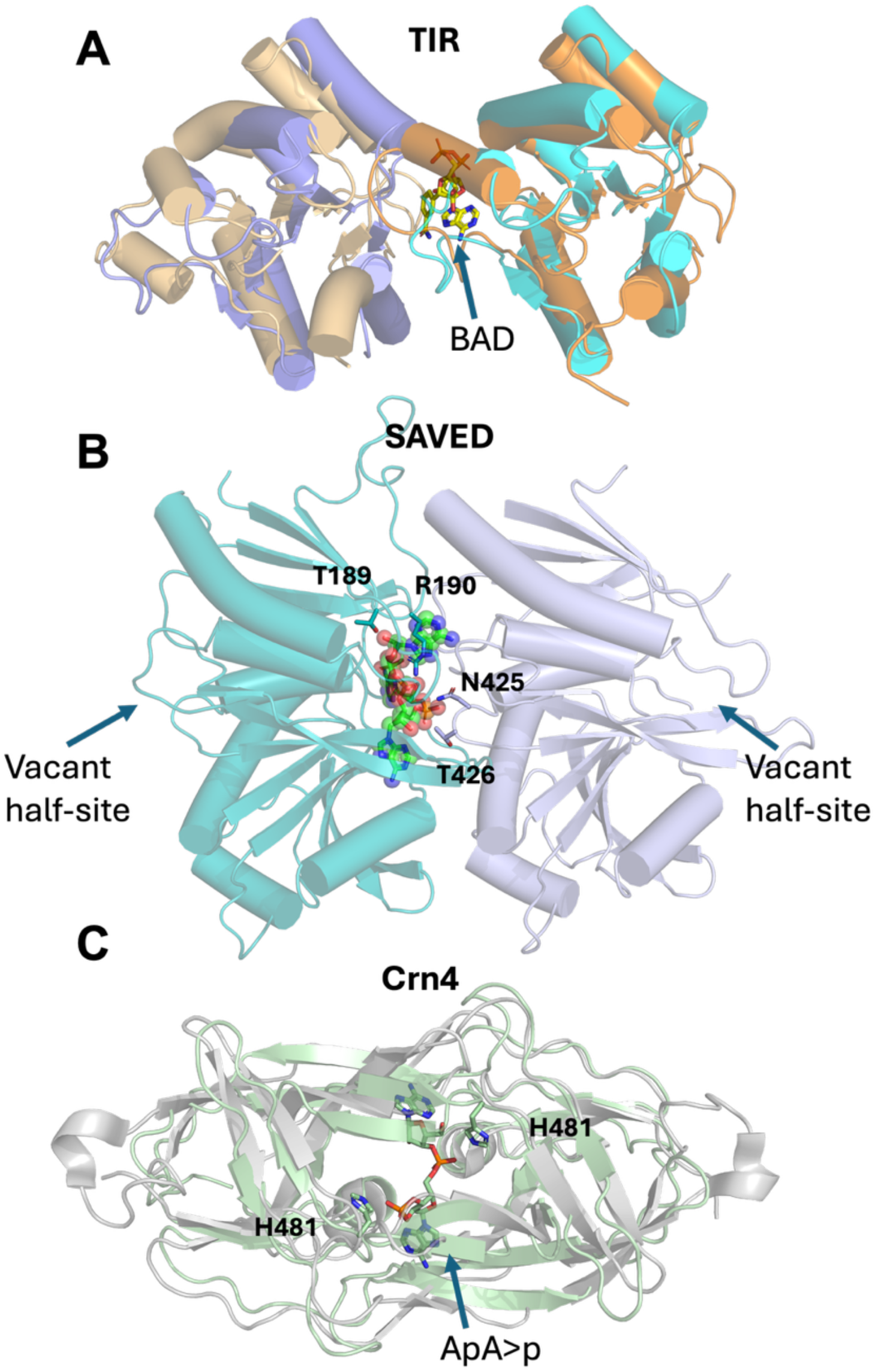
Analysis of TSR1 domains and nucleotide binding sites. **A**. The adjacent TIR domains of the TSR1 complex (orange, gold) are aligned with the TIR domains of the Cat1 effector (blue, light blue) (PDB 9MUO). The NAD+ analogue BAD, which was co-crystallised with Cat1 to locate the NADase active site (7), is shown in stick format. **B**. The SAVED domains of the TSR1 dimer (blue, teal) are shown with cA_3_ sandwiched between them. The cA_3_-interacting residues T189/R190 and N425/T426 are shown in stick format. **C**. The Crn4 domain of the TSR1 protein is shown (green) structurally aligned with the stand-alone Crn4 dimer (PDB 9QS9) (grey). The active site residue H481 is labelled and shown in stick format, along with the ApA>p reaction product.

Examining the structure of the Crn4 domain (Fig. 5C), the cyclic phosphate terminus of the ApA>p product is positioned close to the catalytic histidine (H481). Presumably cA_3_ bound in the Crn4 active site during crystallisation was turned over by the enzyme.

### Investigation of the quaternary structure of TSR1 in the presence and absence of cA_3_ by native mass spectrometry

To characterise the quaternary structure of TSR1 in the presence and absence of the cA_3_ activator, we turned to the technique of native mass spectrometry, which is particularly suitable to determine the oligomeric structure of protein complexes and their ligands as the biomolecules are gently transferred into the gas phase under non denaturing conditions. This technique allows the identification of native-like protein assemblies detected by their mass to charge ratio. We used the H481A variant of TSR1 to avoid turnover of cA_3_ for these experiments. In the absence of cA_3_, TSR1 behaved as a dimer (Fig. 6A), consistent with the hypothesis that the dimeric Crn4 domain prevents dissociation of monomers. At a protein: cA_3_ ratio of 2:1, only two binding sites were observed (Fig. 6B), probably corresponding to the Crn4 and internal SAVED binding sites. At a protein:cA_3_ ratio of 1:3, we observed 3 species corresponding to 1, 2 or 3 bound cA_3_ moieties (Fig. 6C). This is consistent with binding of one cA_3_ molecule in the SAVED dimer, one in the Crn4 dimer and potentially a third in the vacant, outward-facing half-site of one of the SAVED domains (Fig. 5B).

**Figure 6.**
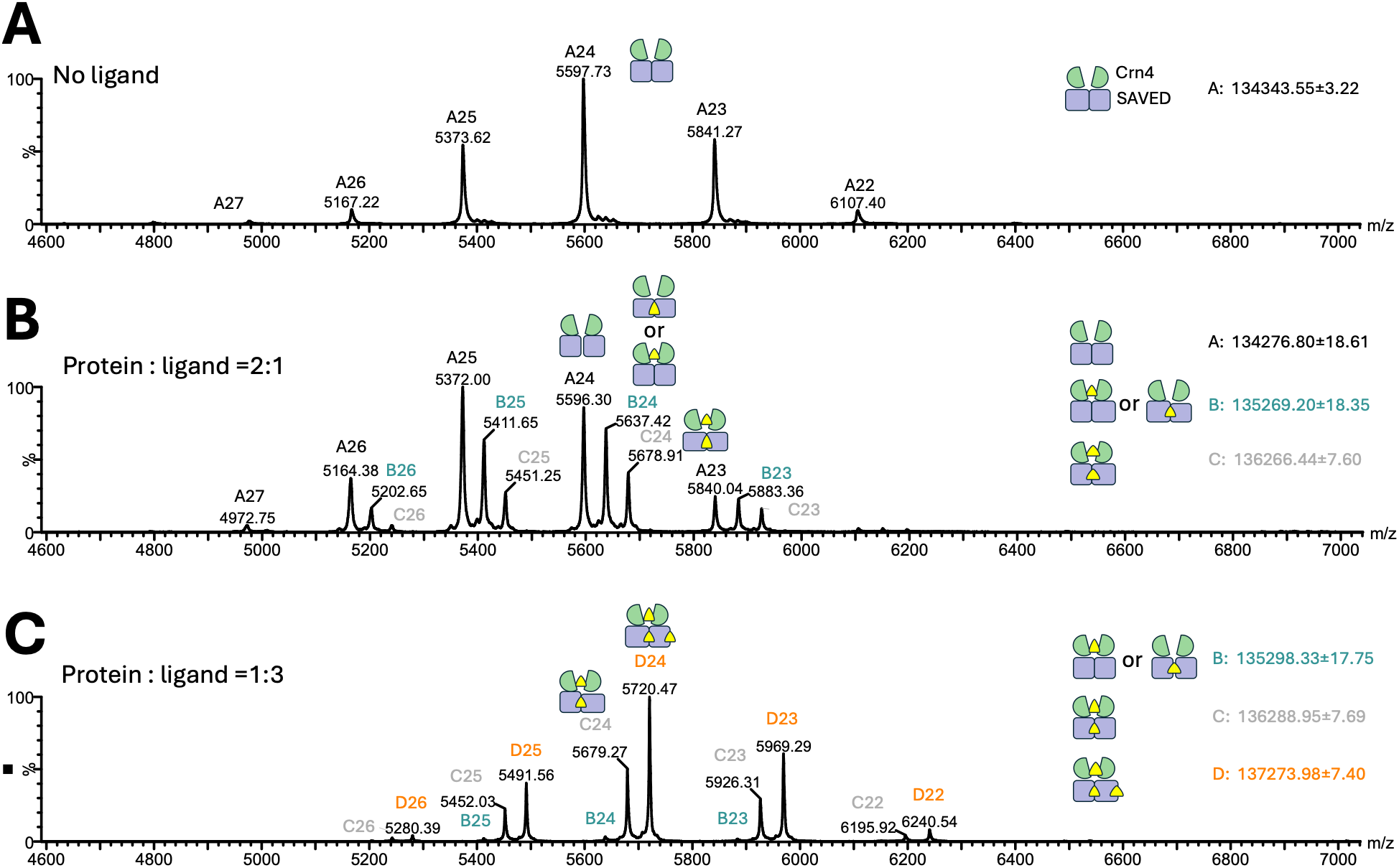
Native Mass Spectrometry of the TSR1 complex. **A**. Native MS spectrum of 5 µM TSR1 dimer (H481A variant) in the absence of cA_3_ shows a dimeric composition. **B**. Native MS spectrum of 5 µM TSR1 dimer in the presence of 2.5 µM cA_3_ revealed 3 complexes: unbound TSR1 (ion series A: 134, 276 Da), TSR1:cA_3_ (ion series B:135, 269 Da), and TSR1:2cA_3_ (ion series C:136, 266 Da). **C**. Native MS spectrum of 5 µM TSR1 dimer in the presence of 15 µM cA_3_ revealed 3 complexes: TSR1:cA_3_ (ion series B: 135, 298 Da), TSR1:2cA_3_ (ion series C:136,288 Da), and TSR1:3cA_3_ (ion series D:137, 273 Da). Full spectra and intact mass spectra are shown in Supplementary fig. S5.

### Removal of the Crn4 domain results in a monomeric, inactive enzyme

The crystal structure of TSR1 demonstrates how the dimeric Crn4 domain bridges two monomers of the protein, stabilising the dimeric organisation. Native mass spectrometry suggested that the dimer is the predominant species even in the absence of the cA_3_ ligand. To investigate this further, we truncated the TSR1 protein at position G465, generating a ΔCrn4 variant. This variant could be expressed and purified to homogeneity (Supplementary fig. 2B). Testing the NADase activity of this variant in the presence of cA_3_ revealed a complete loss of activity (fig. 7A), suggesting that this perturbation disrupted the structure to the extent that an active dimeric TIR domain could no longer be attained. We further analysed the quaternary structure of the variant using analytical SEC, revealing that the ΔCrn4 variant was monomeric in both the presence and absence of cA_3_, unlike the dimeric wild-type protein (fig. 7B).

**Figure 7.**
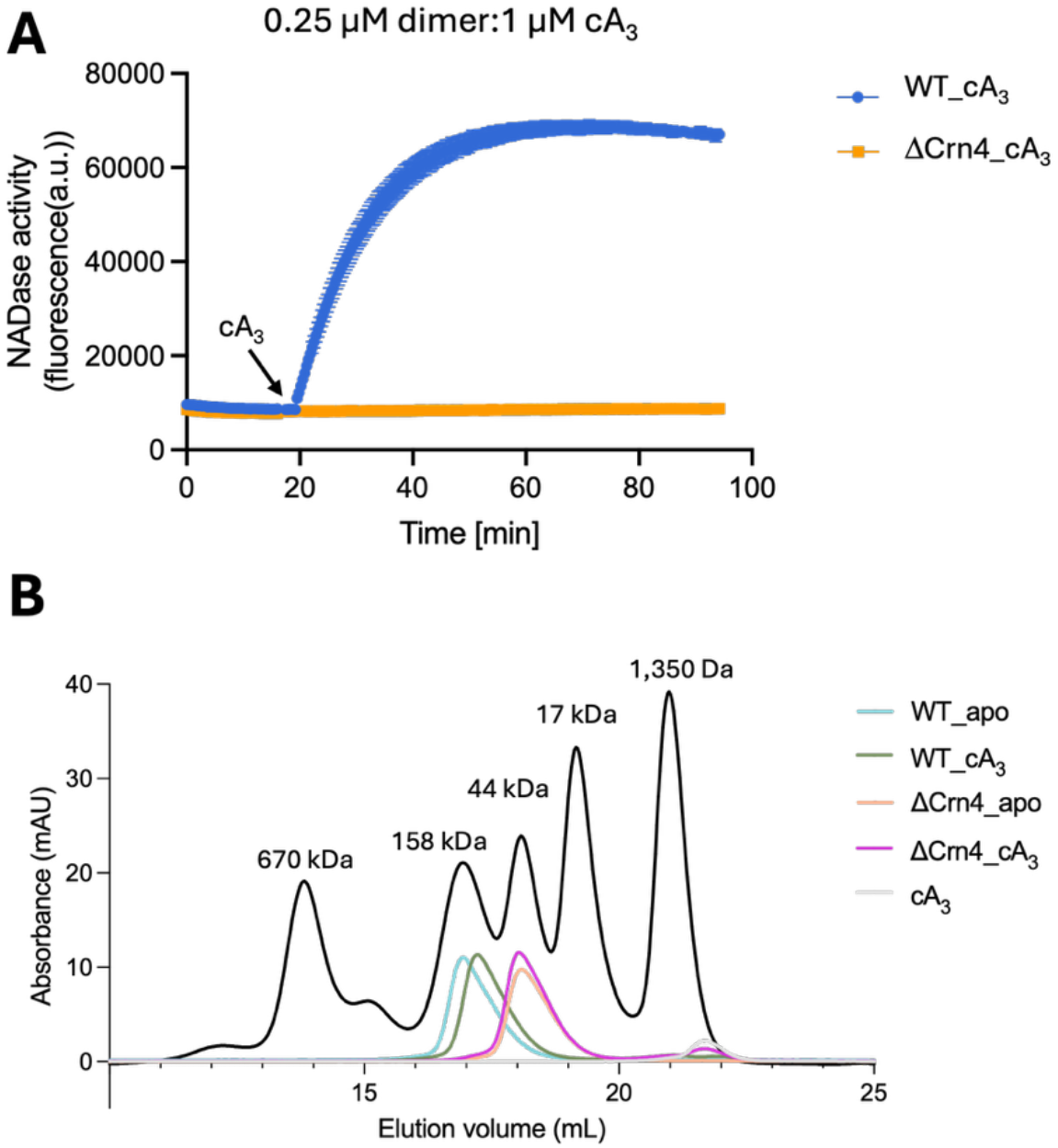
Truncation of TSR1 to remove the Crn4 domain results in an inactive monomer. **A**. NADase activity of WT and ΔCrn4 truncation mutant were analysed by continuous fluorescence assay. Reactions were incubated at 37 °C with the substrate εNAD+ at 500 µM and 0.25 µM TSR1 (WT or ΔCrn4). cA_3_ (0.5 µM) was added to initiate the reaction (arrow). Data points represent the mean ± standard deviation (SD) of three replicate within a single assay. **B**. Analytical size exclusion chromatography (SEC) of WT, and ΔCrn4 TSR1 in the presence and absence of cA_3_. 1 mg/ml of protein in 100 μL was injected into the column. The WT protein eluted in a volume consistent with a dimeric composition, irrespective of the presence of cA_3_. The ΔCrn4 variant eluted later, consistent with a monomeric composition. SEC standards comprised thyroglobulin, γ-globulin, ovalbumin, myoglobin, vitamin B12, with masses indicated above peaks.

## Discussion

TIR domains that degrade NAD+ are one of the most widespread enzymatic effectors found in bacterial antiphage immune systems (13,45), but are rather rare in type III CRISPR systems (16). Typically, TIR-containing effectors are activated by multimerisation driven by recognition of a primary or secondary signal of viral infection (45). The TSR1 effector appears unique in two important respects: it does not change quaternary structure on activation, and it includes a mechanism for its own deactivation. Both of these aspects are worthy of further discussion.

Firstly, our analyses clearly show that TSR1 is always a dimeric protein, regardless of its active state. Despite our best efforts, we could not obtain crystals of the apo form of TSR1. We thus cannot explain the mechanism of activation on cA_3_ binding at a molecular level. We speculate that the apo protein retains a dimeric composition as it is “stapled” by the Crn4 domain, which is a domain-swapped dimer (25). The adjacent SAVED domains may thus be held in close approximation to one another, ready to accept a cA_3_ molecule in the interface, adopt the dimeric head-to-tail conformation that we have crystallised and bring the two TIR domains into an active dimeric state. Although enzymatically active TIR proteins form a shared active site between two adjacent domains, TIR effectors are rarely if ever dimeric in their active form, tending to assemble into larger complexes such as tetramers, octamers and extended filaments (45). Indeed, subunit mixing experiments have demonstrated that the CBASS TIR-SAVED effector is completely inactive as a dimer, relying on filamentation for activation (9). This constraint may represent a mechanism to prevent aberrant activation of TIR immune effectors in the absence of a strong signal of infection, reducing the metabolic burden of these defence systems (9).

Secondly, TSR1 has the ability to auto-deactivate due to the presence of a fused Crn4 domain. Crn4 ring nucleases were recently described as versatile enzymes that degrade a range of cOA species (25). Here Crn4 has a clear specificity for cA_3_ – the first ring nuclease with this property as the standalone Crn4 enzyme shows a preference for cA_4_ (25). We have shown that deactivation of the Crn4 domain results in a more active NADase when TSR1 is activated by cA_3_ *in vitro*. This fits the paradigm that the majority of type III CRISPR effectors are encoded along with an intrinsic or extrinsic ring nuclease enzyme to regulate their activity (24). This in turn supports the hypothesis that type III CRISPR is not generally a system that results in programmed cell death / abortive infection. This is a point of departure from most of the characterised TIR-containing immune systems, which in effect don’t have an “off-switch”.

In conclusion, TSR1 joins NucC (21) and SAVED-CHAT (46) as a rare example of a type III CRISPR effector that is activated by cA_3_. As this cyclic nucleotide is also synthesised by some CBASS systems, such effectors can play a role in multiple antiviral defence pathways. The inclusion of a ring nuclease domain in TSR1 appears to be an adaptation to a function in CRISPR defence, which generally favours a means to limit effector activity. A side effect of this fusion may have been to force the enzyme into an obligate dimeric structure, in contrast to the filament form of the closely related CBASS effector TIR-SAVED. TSR1 may be the first clear example of an active dimeric TIR effector in immune defence, supporting the hypothesis that TIR multimerization beyond the dimer is a regulatory adaptation.

## Data Availability

The protein structure coordinates and data have been deposited in the Protein Data Bank with deposition code 31JM.

## Supplementary Data

Supplementary data are available at NAR Online.

## Author Contributions

YS: data acquisition, analysis and interpretation. SM: crystallographic data acquisition, analysis and interpretation; SS: native MS data collection and analysis; MFW: funding acquisition, conception, data interpretation. All authors contributed to the drafting and revision of the manuscript.

## Acknowledgements

Thanks to Carlos Penedo and Sally Shirran for helpful discussions and to the British Mass Spectrometry Society for their research support grant. We acknowledge Diamond Light Source for time on Beamline i04 under proposal number mx 41060.

