## Supplemental Data for "A self-limiting dimeric TIR effector specialised for type III CRISPR-mediated immunity"

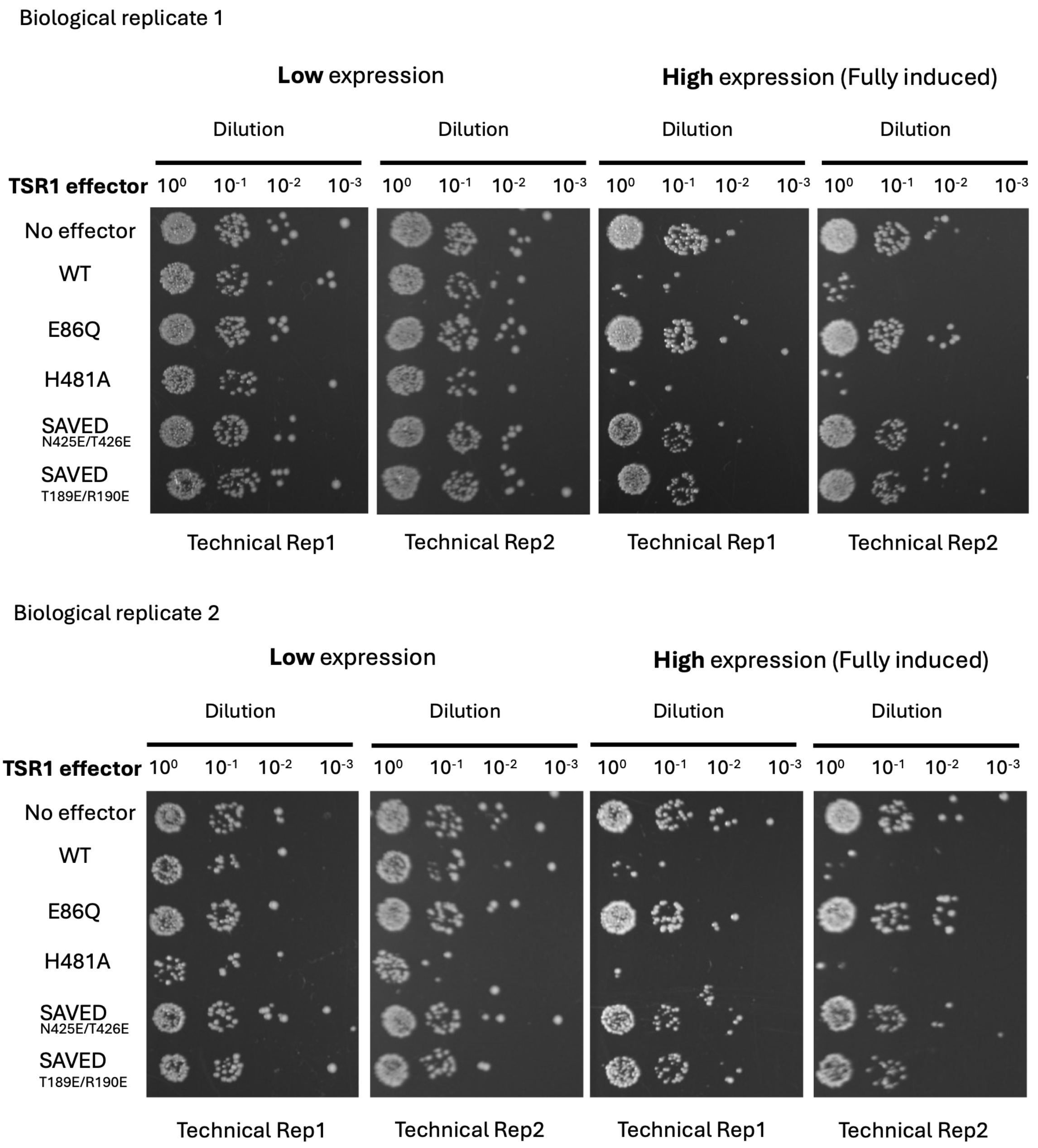

**Supplementary Figure 1.** Plasmid challenge assay under low (no induction) and high level (0.2 % w/v D-lactose and 0.2 % w/v L-arabinose) induction of TSR1 WT and variants, including E86Q, H481A, $SAVED^{N425E/T426E}$, $SAVED^{T189E/R190E}$. Representative plates of two biological replicates with two technical replicates with dilution factor from $10^0$ to $10^{-3}$ are shown.

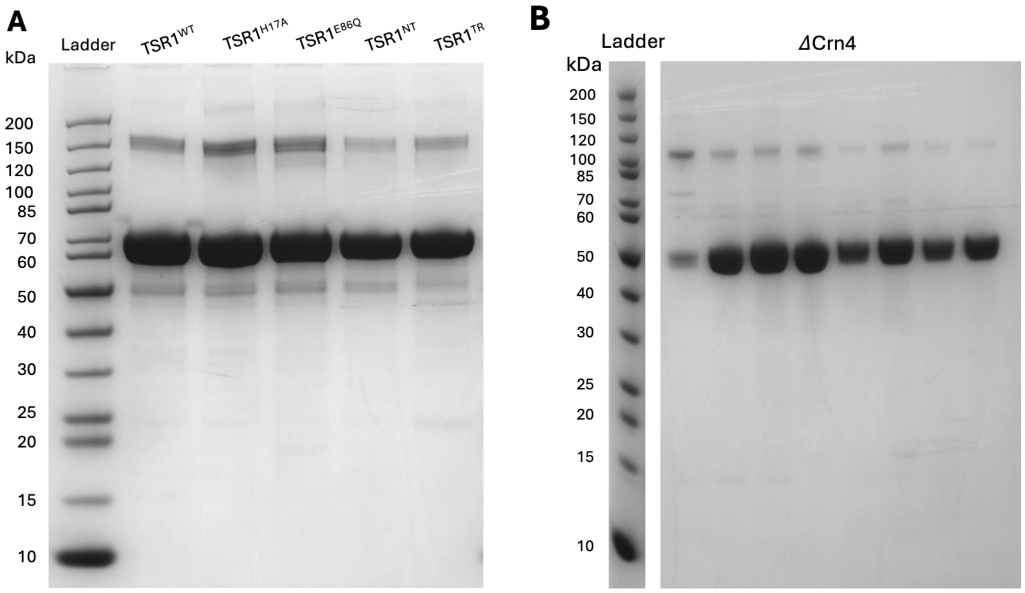

**Supplementary Figure 2. SDS-PAGE of purified TSR1 and variants used in assays** (A)SDS-PAGE of purified TSR1 and variants used in assays, showing the indicated monomeric and dimeric species.10 µg protein of wild type TSR1 and variants were analysed by SDS-PAGE and Coomassie staining. (B) Purified *Δ*Crn4 from SEC was analysed by SDS-PAGE and Coomassie staining.

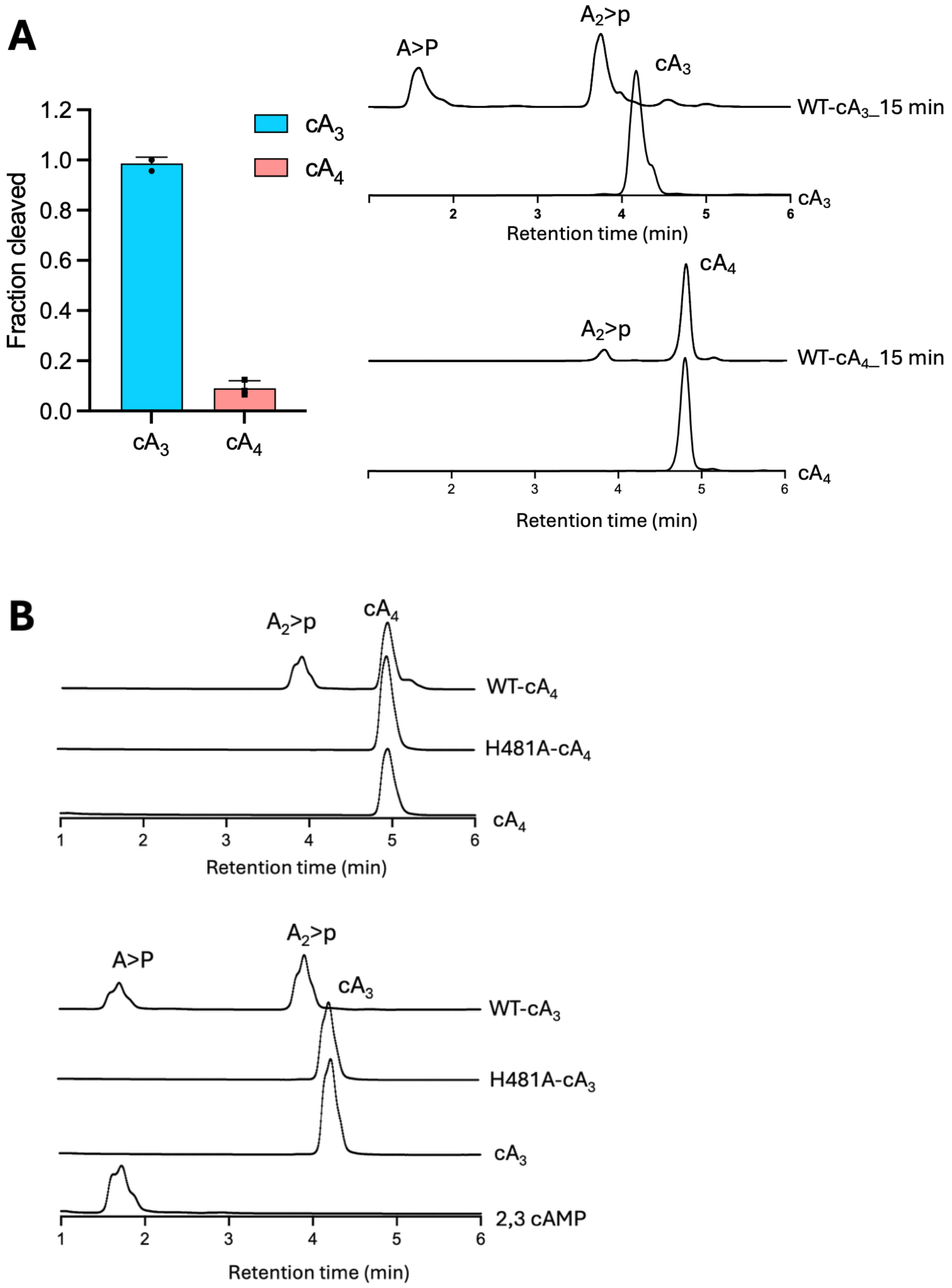

**Supplementary Figure 3. A. TSR1 exhibits ring nuclease activity with a preference for $cA_3$.** Reactions were incubated at 37 °C for 15 min with the indicated cOA species (50 µM) and WT TSR1 (0.5 µM dimer). $cA_3$ and $cA_4$ standards served as positive controls. Peaks of substrate and cleavage products were quantified and data plotted as fraction cleaved over 15 min. Data points represent the means of triplicate experiments with standard deviation shown. Source data are provided in the Source Data file. **B.** HPLC chromatograms of $cA_3$ and $cA_4$ cleavage reactions. 50 µM $cA_3$ or $cA_4$ was incubated with WT or H481A separately at 37 °C for 1 hour. Standards of $cA_3$, $cA_4$ and 2,3 cAMP serve as controls.

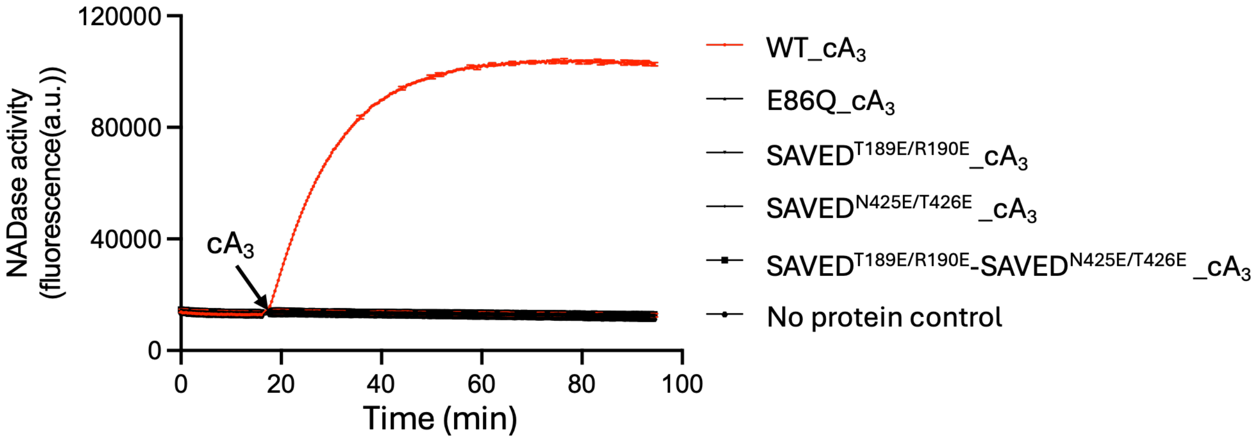

**Supplementary Figure 4. cA₃ binding at SAVED domain is essential for activation of the TIR domain**. NADase activity of WT and variants were analysed by continuous fluorescence assay. Reactions were incubated at 37 °C with the substrate εNAD+ at 500 μM and protein dimer at 0.25 μM. 0.5 μM $cA_3$ was added at cycle 50 (16 min). The data are means plotted with the standard deviation for triplicate experiments.

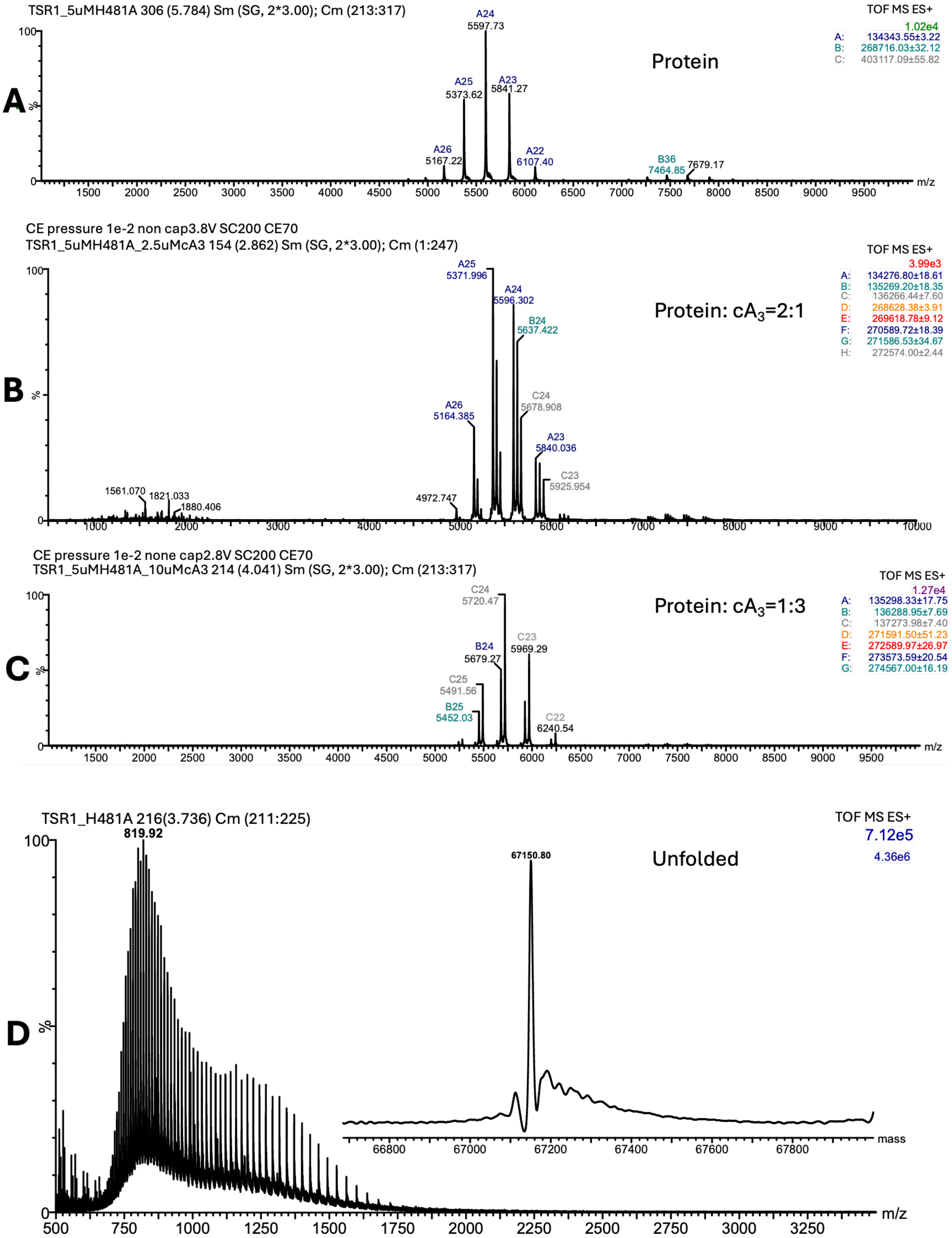

**Supplementary Figure 5. Full spectrum Native MS results for TSR1 H481A. A.** Full Native MS spectrum of 5 μM TSR1H481A dimer. Ion series A, B, C represent 3 different complexes, unbound TSR1 dimer, tetramer, and pentamer complexes that were not further analysed in this study. **B.** Full Native MS spectrum of 5 μM TSR1 H481A dimer in the presence of 2.5 μM $cA_3$. Ion series A, B, C correspond to 3 different complexes, unbound TSR1, TSR1:1$cA_3$, TSR1:2$cA_3$. Ion series D–H are substantially less abundant than A–C and correspond to higher-order tetrameric complexes that were not further analysed in this study. **C.** Full Native MS spectrum of 5 μM TSR1H481A dimer in the presence of 15 μM $cA_3$. Ion series A, B, C represent 3 different complexes, TSR1:1$cA_3$, TSR1:2$cA_3$, TSR1:3$cA_3$. Ion series D–G correspond to higher-order tetrameric complexes that were not further analysed in this study. All samples were sprayed from 150 mM ammonium acetate. **D.** Intact Mass measurement under unfolded conditions obtained from purified Apo TSR1H481A, showing the mass at 67150 Da.

### Table S1. Oligos and synthetic gene sequences used in this study.

| Oligos | 5' to 3' |
|---|---|
| saved-T189R190EE-Fwd | TCGGCTTACAAGAGGAACCTGAGCCTAACCACTTG |
| saved-T189R190EE-Rev | AGGCTCAGGTTCCTCTTGTAAGCCGATTTCAATTTC |
| saved-N425T426EE-Fwd | GCTGGTGCGAAGAGCTTACTATTGACTTCTATG |
| saved-N425T426EE-Rev | TCAATAGTAAGCTCTTCGCACCAGCGGGCAGTAAG |
| DeltaCrn4-Fwd | TTCCCACAG**T**GAGTACCCCAGGTGGACG |
| DeltaCrn4-Rev | ACCTGGGGTACTC**A**CTGTGGGAAGACAC |
| TirE86Q-Fwd | TACCGAAAACC**C**AAGCTCCCCTTTTAG |
| TirE86Q-Rev | AGGGGAGCTTGG**G**TTTTCGGTACGAATG |
| Crn4H481A-Fwd | ACTTAACACCG**GCA**GATGTGACCTACTAC |
| Crn4 H481A-Rev | TAGGTCACATC**TGC**CGGTGTTAAGTTAAT |
| Gblock_*tsr1* | GCGCCCATGGCACATATGACATCGAGCCGTCCGCCAGGTCGTGGCAACGGTCCTGTCTTCATTTCATA<br>TCACCAAAAATCTGGAGCAGCTGACGCGGAATTTATTGAGACTTATTTACGTGCAGGCGGTATCGTACCT<br>TGGCGTGATATTCGTGATTTAGAAGCGGGCACGGTAGAGCGCAACATTACTCAGGCGTTTGAGGAAGGA<br>TTATCTGGCGGGGTACTGTTACTTAGTTCCGGTATTAGCGAAAGCTCATTCGTACCGAAAACCGAAGCTC<br>CCCTTTTAGTGGGTGCCCACAAGGCAGATCCCGAGGGGTTTCAGTTACACATCATTAATACATTTCGCAA<br>ACCGGGATCTCCGGATGAGTGCGATTTTAAGGCCCCCGGAAAGCAATTGAAGACCAAGTACCCGGAA<br>GCCGAGCAATTAAACGATCATTTGCAGCGTCGCCTGCTGCACTCAGACGACAAAGGGGGAAAGCCCG<br>TGTCGGAGCTTAATTTAGTACTTCGCGATCTGTTGCGCAATCGTTTGAAAGTACGCCGTCCCCAATTGGG<br>AGACGGTGAAATTGAAATCGGCTTACAAACGCGTCCTGAGCCTAACCACTTGCCGGCGGACGGGCGT<br>ACGGTCCCGGAGGCAGATTTACACATTCGCTTGCGCCAGGACGCGGCTACGCAAATTCCAGAGGAGC<br>TGGATTATCGCTGTTTGCAGCAAGCATTGCCAGTGCTTATCGATGAGCTTCACGCGGCTCGCATTCGTC<br>GCGTGTTGTTCCGTGGGGGTTGTCATCCCAGTTTGGCGTGGGCATTGGGGGTAGCACTTCCGCACGCG<br>CGTGAGATCGAGCACTTCACTTGGCGTGACACCTACGGCAAGGACTGGGCATCAGCGGATGAACCTG<br>CTGAGCGTTCAACGAGCATTCATTTAGAGACACTTAATCCGGACGGCTCACGTCGTGCTTTGGGGTTCG<br>CCCCTGACGAGATTCCGAGTGGGGCGGAGTTACGTCGTGCTTTATGCGGAGATGCCCCGGCCAAGAA<br>CGCTGTAGTCTTACTTGCGGCTGATGACCTGCGCCCACAGCCTCTGTTGGCCCTTGCTGAGAAGTTGG<br>ATGATTCACCGGTCCTGGTAATTAATTTGCACACCCCGAGCGCTGACGGAGCTAAAAAGTGGATCGACC<br>ATGCCGAGGGTGCCGGGTTAGCCCGTCGCGTGGGTGAGATTCTTCGCCGTTTGCGCGATTTGGCTAAA<br>TTACATCTTGCTATTTCGGCCCCTGCTGCTATGGCGGCTCTTACTGCCCGCTGGTGCAATACTCTTACTA<br>TTGACTTCTATGAGCTTGGAAACACCGGTATGGGTGCCCGCGAATACATTCGTGTGTTACGTACCGAAA<br>GCGGAAACAAATCTCCCATTACCGGTGTCTTCCCACAGGGAGTACCCCAGGTGGACGAAGTCCGTAAA<br>TTAATTAACTTAACACCGCATGATGTGACCTACTACCCAGAAGCTGGGGAACCATTTACATGGGCAGCA<br>CCCGAAGGTCCAGACCAGTGGGTGCGTCGCCAGGAGCAGTCAGAGGAGCTTCCGTCGTTGCGTGTC<br>CAAGGTTTGGAAATTCCAGTTACCCGTATCCGTCAGGGTACCATCGCACCTGTGCCTGACCCGATGCC<br>TGGTGTCGGTTACATTGTTCCCCGTATTAGCGCGGAAACCGCGCGCCGTCCGGACTTTTTCTTCCCTCA<br>CGGCGAAGTACGCGGACAGGGCGGAGGAATCATCGGTTGTCGCCGCCTGGGGTGTTTCGAAGCGGT<br>TAGTAATCGTGTACGCCCTTATTTGGAGTTGCTTGACCCGGTGCCTCAAGATTGACTCGAGGGATCCCG<br>CG |

**Table S2. Data collection and refinement statistics for the structure of TSR1 in complex with $cA_3$**

| | Apr TSR1 |
|---|---|
| **Data processing** | |
| Space group | P $2_1$ $2_1$ $2_1$ |
| Cell dimensions | |
| a, b, c (Å) | 125.2, 150.9, 245.5 |
| α, β, γ (°) | 90, 90, 90 |
| Resolution (Å) | 111.50 – 2.68 |
| | (2.73 – 2.68) |
| $R_{merge}$ | 0.129 (4.360) |
| $I/\sigma(I)$ | 12.0 (0.3) |
| Completeness (%) | 99.9 (97.2) |
| Average redundancy | 13.8 (14.3) |
| $CC_{1/2}$ | 0.999 (0.300) |
| $V_m$ ($Å^3$/Da) | 2.91 |
| Solvent (%) | 57.7 |
| **Refinement** | |
| Unique reflections | 130441 (6291) |
| $R_{work}$ / $R_{free}$ | 23.9 / 28.4 |
| Geometric deviations | |
| Bonds (Å) / Angles (°) | 0.003 / 0.645 |
| No. atoms (non H) | |
| Protein | 24584 |
| Water | 75 |
| cA3 | 198 |
| ApA>p | 176 |
| **B factors ($Å^2$)** | |
| Protein | 103.0 |
| Water | 80.5 |
| cA3 | 86.2 |
| ApA>p | 108.4 |
| Ramachandran outlier (%) | 0.03 |
| Rotamer outliers | 0.87 |
| Molprobity score / centile (%) | 1.53 / 100 |
| PDB Code | 31JM |
